# NEK1 haploinsufficiency impairs ciliogenesis in human iPSC-derived motoneurons and brain organoids

**DOI:** 10.1101/2024.02.29.582696

**Authors:** Sorce Marta Nice, Invernizzi Sabrina, Casiraghi Valeria, Santangelo Serena, Lattuada Chiara, Podini Paola, Brusati Alberto, Silva Alessio, Peverelli Silvia, Quattrini Angelo, Silani Vincenzo, Bossolasco Patrizia, Ratti Antonia

## Abstract

Primary cilia are microtubule-based organelles acting as specialized signalling antennae that respond to specific stimuli to maintain cellular integrity and homeostasis. Recent studies indicate defective primary cilia in post-mortem human brains and animal models of neurodegenerative conditions, including Amyotrophic Lateral Sclerosis (ALS). Heterozygous loss-of-function mutations (LOF) in *NEK1* gene are present in about 1% of familial and sporadic ALS cases. The protein kinase NEK1 regulates various cellular processes, including ciliogenesis, but a clear link between *NEK1* LOF mutation in ALS and primary cilia is unknown. In this study we generated a human iPSC line carrying a *NEK1* LOF mutation by gene editing, leading to NEK1 protein haploinsufficiency. In differentiated iPSC-motoneurons (MNs) we observed that primary cilia were significantly shorter in *NEK1*-LOF iPSC-MNs compared to wild-type (WT) iPSC-MNs and that also the percentage of ciliated iPSC-MNs was significantly decreased in *NEK1*-LOF cells. We also investigated ciliogenesis in *NEK1*-LOF iPSC-brain organoids confirming that primary cilia were thinner with no apparent alteration in the ultrastructure by transmission electron microscopy.

Our data suggest that NEK1 protein plays a role in regulating ciliogenesis in both 2D and 3D human iPSC-derived neuronal models and that *NEK1* LOF mutations associated to ALS, leading to *NEK1* haploinsufficiency and likely to reduced kinase activity, impair primary cilium formation. The involvement of ciliogenesis dysfunction in ALS deserves further investigation providing novel therapeutic targets and strategies to be addressed for this incurable disease.

## 1. INTRODUCTION

Primary cilia are microtubule-based organelles present in most cell types and serve as specialized “signalling antennae” capable of responding to specific stimuli and regulating cellular integrity and homeostasis (1). In neuronal cells, they contribute to essential functions such as mechanosensing and signal transduction mediated by Sonic Hedgehog, Wnt, and G Protein-Coupled receptors (GPCR), which favour adaptive responses to environmental changes (2,3). A group of human hereditary disorders, collectively known as ciliopathies, are caused by mutations in several genes regulating primary cilia biogenesis, structure and functioning and affecting multiple organs and body systems (1). Moreover, emerging studies suggest that defective primary cilia are present also in human post-mortem brains and animal models of various neurodegenerative conditions including Parkinson and Alzheimer diseases and Amyotrophic Lateral Sclerosis (ALS) (4). In particular, in ALS, transgenic SOD1 G93A mice exhibit a decreased number of ciliated motoneurons in the spinal cord during disease progression (5). Interestingly, another ALS causative gene, the NIMA-related kinase *NEK1*, is involved in ciliogenesis (6). Indeed, recessive *NEK1* mutations cause skeletal ciliopathies such as Short-rib thoracic dysplasia (SRTD) (7,8) and Axial spondylometaphyseal dysplasia (SMDAX) (9). Dominant *NEK1* loss-of-function (LOF) mutations are instead associated with about 1% of both familial and sporadic ALS cases (10–12), but the mechanistic link between *NEK1* haploinsufficiency and ciliogenesis in ALS is unknown. Over the past decade the development of both two-dimensional (2D) and three-dimensional (3D) *in vitro* systems derived from induced Pluripotent Stem Cells (iPSCs) has greatly favoured the establishment of human neural disease models to investigate pathomechanisms related to neurodegenerative disorders while maintaining the patients’ genetic background. In this view, iPSC-motoneurons (iPSC-MNs) with heterozygous *NEK1* LOF mutations associated with ALS were recently generated to study DNA damage response and repair, microtubule homeostasis and nuclear import (13,14), but the impact of such *NEK1* gene mutations on primary cilia formation and functioning has never been investigated so far.

## 2. MATERIALS AND METHODS

### 2.1 iPSC gene editing and characterization

iPSCs from a healthy donor were already available and obtained upon reprogramming of primary fibroblasts with CytoTune®-iPS 2.0 Sendai Reprogramming Kit (Thermo Fisher Scientific) after receiving written informed consent (Ethic committee approval n. 2015-03-31-07) as previously described (15). *NEK1* gene editing was performed using the CRISPR/Cas9 system with the recombinant Alt-R HiFi Cas9 Nuclease, the single-guide RNA and Alt-R HDR Donor oligo (IDT). Sanger sequencing confirmed the insertion of the CTATA nucleotides causing a frameshift and a premature stop codon (ins(CTATA)_Arg261Profs*19) in heterozygous state. The *NEK1* gene-edited iPSC karyotype was assessed by Q-banding. Characterization of stemness and pluripotency markers was conducted by Real-time PCR (Supplementary Table 1 for primer pairs) and immunofluorescence (Supplementary Table 2 for antibody conditions), respectively, as already described (15). NEK1 protein content was evaluated by Western blot analysis with specific antibodies (Supplementary Table 2).

### 2.2 iPSC-motoneurons and iPSC-brain organoids differentiation

iPSC-motoneurons (iPSC-MNs) were obtained as previously described (16). Briefly, iPSCs were seeded in poly-HEMA (poly-2-hydroxyethyl methacrylate)-coated dishes (Merck) for embryoid bodies (EBs) formation and cultured in different media supplemented with specific factors. After 17 days, EBs were dissociated and cells were plated on poly-D-lysine/laminin-coated dishes (Merck) and cultured in neural differentiation medium for 30 days.

For the generation of human brain organoids, iPSCs were initially placed in ultra-low attachment plates for 48 hours to obtain spherical floating colonies. Subsequently, organoids were cultured for a duration of 60 days on a shaker with the addition of specific neuronal differentiation factors as described (17).

### 2.3 Immunofluorescence and image analysis

iPSCs, iPSC-MNs and iPSC-brain organoids were fixed with 4% paraformaldehyde for 20 minutes at room temperature (RT). iPSCs and iPSC-MNs were permeabilized and incubated with specific antibodies as previously described (16)(all antibodies used are listed in Supplementary Table 2).

Whole iPSC-brain organoids were permeabilized with 0.5% Triton X-100 in blocking solution with 2% Bovine Serum Albumin (BSA; Merck) in PBS for 3 hours at RT. Primary antibodies were incubated at 4°C O/N in blocking solution and then the secondary fluorescent antibodies for 8 hours at RT (Supplementary Table 2)(18).

Images were acquired using the Confocal Eclipse Ti2 microscope (Nikon) at 60x magnification as Z-stacks (0.2 μm step size) for iPSCs and iPSC-MNs and at 10x magnification for whole iPSC-brain organoids with 0.5 μm step size. For image analysis, at least 100 cells per sample were considered for each biological replicate. To measure cilia length, the CiliaQ plugin of the ImageJ software was used (19) and obtained values from each replicate were pooled.

### 2.4 Morphologic analysis

iPSC-brain organoids were fixed in 0.12 M phosphate buffer containing 2% glutaraldehyde followed by osmium tetroxide. After dehydration in a graded series of ethanol preparations, organoids were cleared in propylene oxide, embedded in Epon. Ultra-thin (60-100 nm) sections were cut with a Leica EM UC6 ultramicrotome, counterstained with uranyl acetate and lead citrate, and examined with a transmission electron microscope (Talos 120C Fei). Images were acquired with a 4kx4k Ceta CMOS camera (Thermo Fisher Scientific).

### 2.5 Statistical analyses

Statistical analyses were performed using GraphPad Prism 9 software by applying Student’s t-test. Results were considered statistically significant if *p* < 0.05.

## 3. RESULTS

### 3.1 Generation of human *NEK1* Loss-of-function (LOF) iPSCs by gene editing

In order to study the impact of *NEK1* mutations on primary cilia formation in ALS, we mimicked *NEK1* haploinsufficiency in human iPSCs by gene editing. We used CRISPR/Cas9 technology to introduce a LOF mutation into a healthy control iPSC line already generated in our lab and fully characterized (15). The 5-nucleotide insertion in the *NEK1* gene resulted in a frameshift leading to a premature termination codon (ins(CTATA)Arg261Profs*19) in heterozygous state as determined by Sanger sequencing (Fig.1A). The gene-edited *NEK1*-LOF iPSCs had a normal karyotype and lacked any gross chromosomal rearrangement (Fig.1B). We confirmed the maintenance of the stemness state of the gene-edited *NEK1*-LOF iPSC line by Q-PCR for the expression of *OCT3/4, SOX2* and *NANOG* genes (Fig.1C) and by immunofluorescence staining for the specific pluripotency markers TRA-1-60, AP (Alkaline Phosphatase) and SSEA-4 (Fig.1D).

**Fig. 1.**
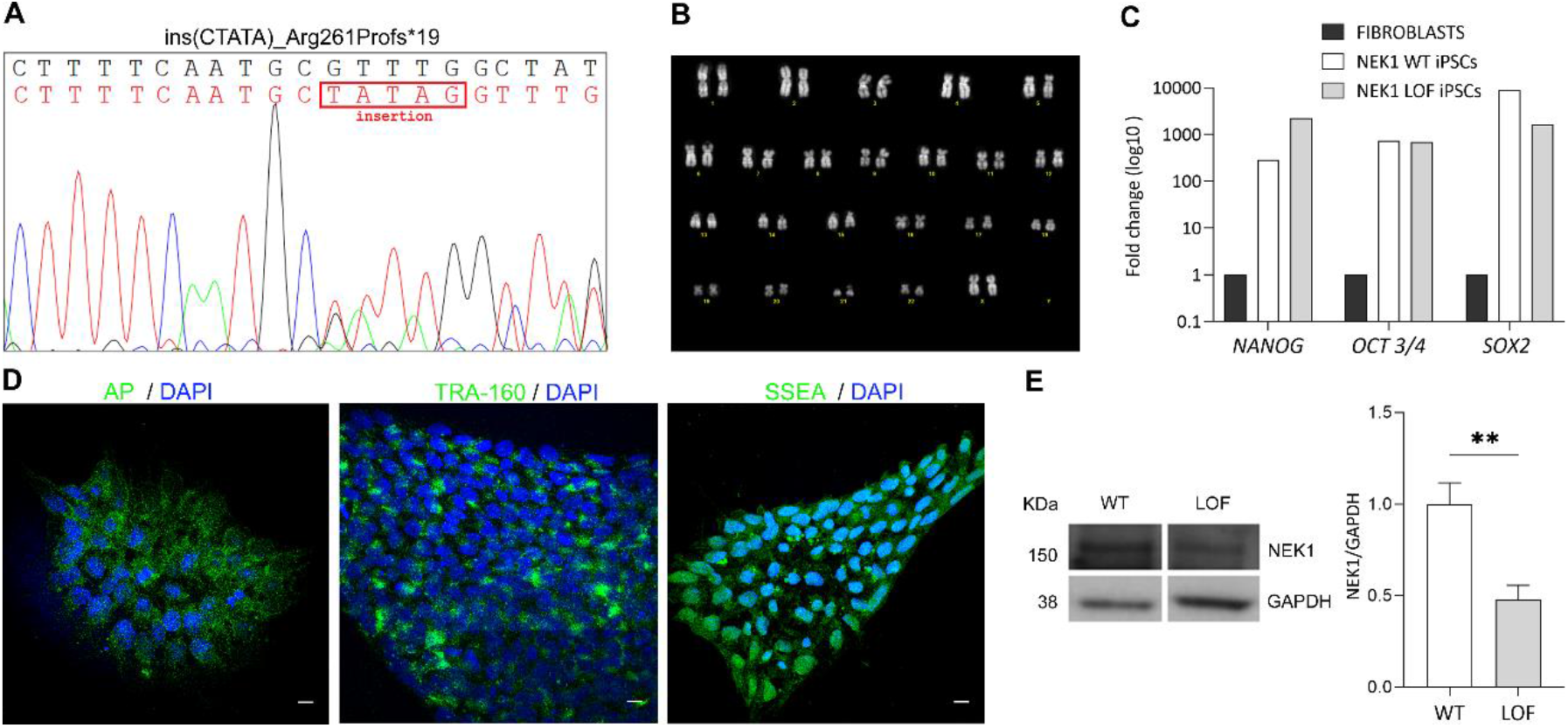
Generation and characterization of human *NEK1*-LOF iPSCs. (A) Sanger sequencing confirmed the presence of the CTATA insertion leading to a premature stop codon (ins(CTATA)_Arg261Profs*19). (B) Karyotype analysis of the NEK1-LOF iPSC line by G-banding. (C) Gene expression analysis of *NANOG, OCT 3/4* and *SOX2* in control fibroblasts, *NEK1*-WT iPSCs and *NEK1*-LOF iPSCs by Real-time PCR. Data were normalized to control fibroblasts. (D) Representative confocal images of pluripotency markers AP, TRA-1-60 and SSEA; Scale bar 10 μm. (E) Representative Western Blot images and densitometric analysis of NEK1 protein (n=3, mean ± SD; Student’s t-test; **p < 0.001).

*NEK1* haploinsufficiency was assessed by Western blot analysis which showed a significant reduction of NEK1 protein level in the gene-edited *NEK1*-LOF iPSCs (0.47X) compared to the wild-type (WT) isogenic iPSC line (Fig.1E).

### 3.2 Human *NEK1*-LOF iPSC-derived motoneurons show defective cilia

To assess the potential impact of *NEK1* haploinsufficiency in the context of ALS disease, we differentiated *NEK1*-LOF and *NEK1*-WT iPSCs into motoneurons (iPSC-MNs) which were confirmed to express neuronal (βIII-tubulin and SMI-312) and motoneuronal (ChAT) markers (Fig.2A). Given the limited understanding of the link between *NEK1* and ciliogenesis in neuronal cells, we investigated primary cilia formation and distribution by immunofluorescence analysis using the acetylated tubulin (ACIII) marker (Fig.2B). A significantly smaller fraction of *NEK1*-LOF iPSC-MNs (36.3%) showed ACIII-positive cilia compared to *NEK1*-WT iPSC-MNs (52.4%) (Fig.3C). Furthermore, *NEK1* haploinsufficiency caused a significant reduction in cilia length in *NEK1*-LOF iPSC-MNs (0.91 µm) in comparison to *NEK1*-WT iPSC-MNs (1.49 µm) as assessed by quantitative image analysis (Fig.4D).

**Fig. 2.**
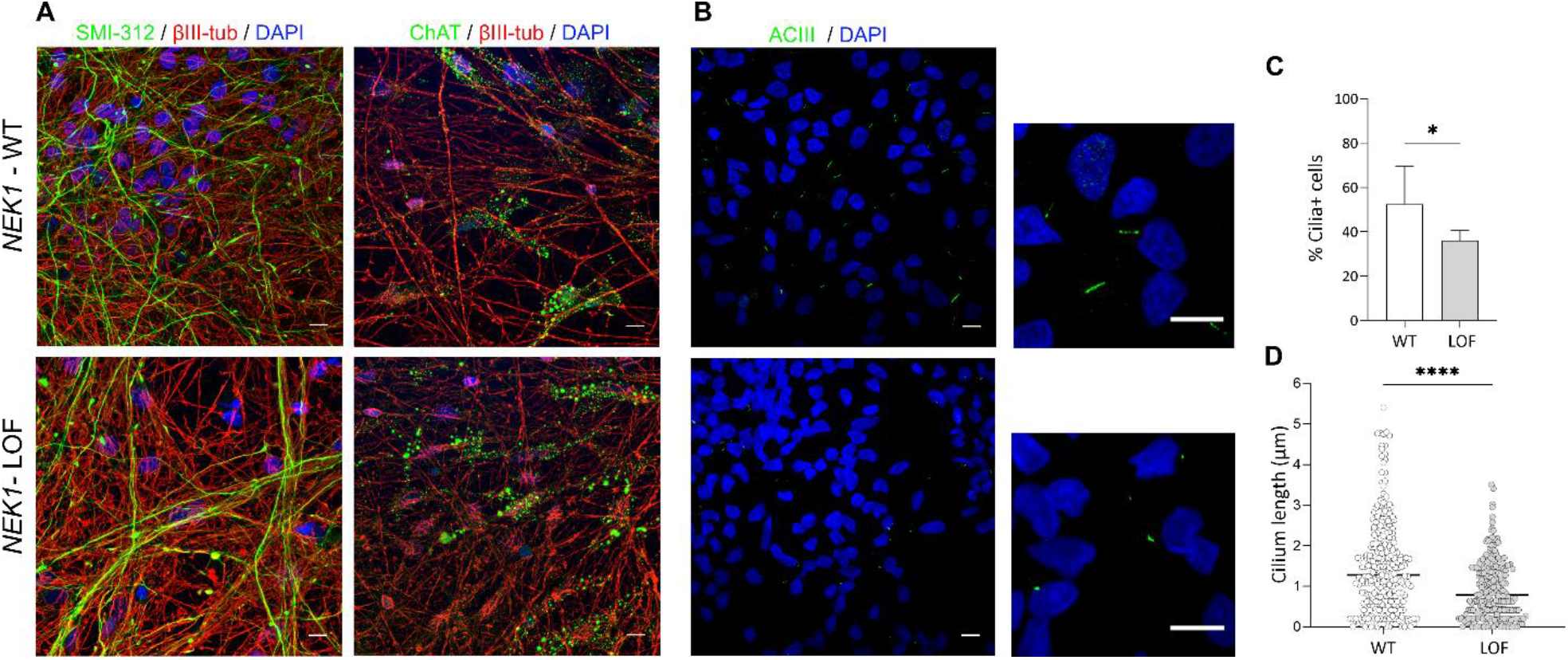
Analysis of primary cilia in human iPSC-derived motoneurons. (A) Representative confocal images of the neuronal markers SMI312 (green) and βIII tubulin (red) and motoneuronal marker ChAT (green) in *NEK1*-WT and *NEK1*-LOF iPSC-MNs. Bar, 10 μm. (B) Representative confocal images of Adenylate Cyclase marker (ACIII) (green) in *NEK1*-WT and *NEK1*-LOF iPSC-MNs. White boxes indicate the area of the adjacent enlarged image; scale bar 10 μm. (C) Quantification of cilium-positive cells and (D) cilium length in *NEK1*-WT and *NEK1*-LOF iPSC-MNs (at least 100 cells counted/sample; n=3, mean ±SD; Student’s t-test; *p<0.05; **** p<0.0001).

### 3.3 *NEK1* LOF mutation impairs cilia size in human iPSC-brain organoids

To further investigate whether *NEK1* has an impact on ciliogenesis in iPSC-derived 3D models, we obtained brain organoids from *NEK1*-LOF and *NEK1*-WT iPSCs following a validated differentiation protocol as described in Materials and Methods (Fig.3A). Organoids were maintained in culture for a maturation time of 60 days (Fig.3B) and subsequently characterized by whole-mount immunofluorescence. We assessed the presence of the neuronal markers βIII-tubulin, MAP2, doublecortin and NeuN along with the cholinergic marker ChAT in both *NEK1*-LOF and *NEK1*-WT iPSC-brain organoids (Fig.3C). Furthermore, iPSC-brain organoids exhibited positivity for the dopaminergic marker TH (tyrosine hydroxylase) and the astrocytic marker GFAP, confirming the presence of a mixed neuro-glial cell population (Fig.3C). We also evaluated the expression of Ki67, a cell proliferation marker of the initial phase of neurogenesis (Fig.3C). No qualitative differences in the distribution of these markers were observed between the two different iPSC-brain organoids. However, by conducting an ultrastructural analysis by transmission electron microscopy (TEM), we observed the presence of thinner and shorter cilia in *NEK1*-LOF iPSC-brain organoids compared to *NEK1*-WT ones as observed in longitudinal sections (Fig.3D). Transversal images at the level of cilium basal body showed no apparent alterations in the 9+0 architecture of the transition fibers (Fig.3E-F).

**Fig. 3.**
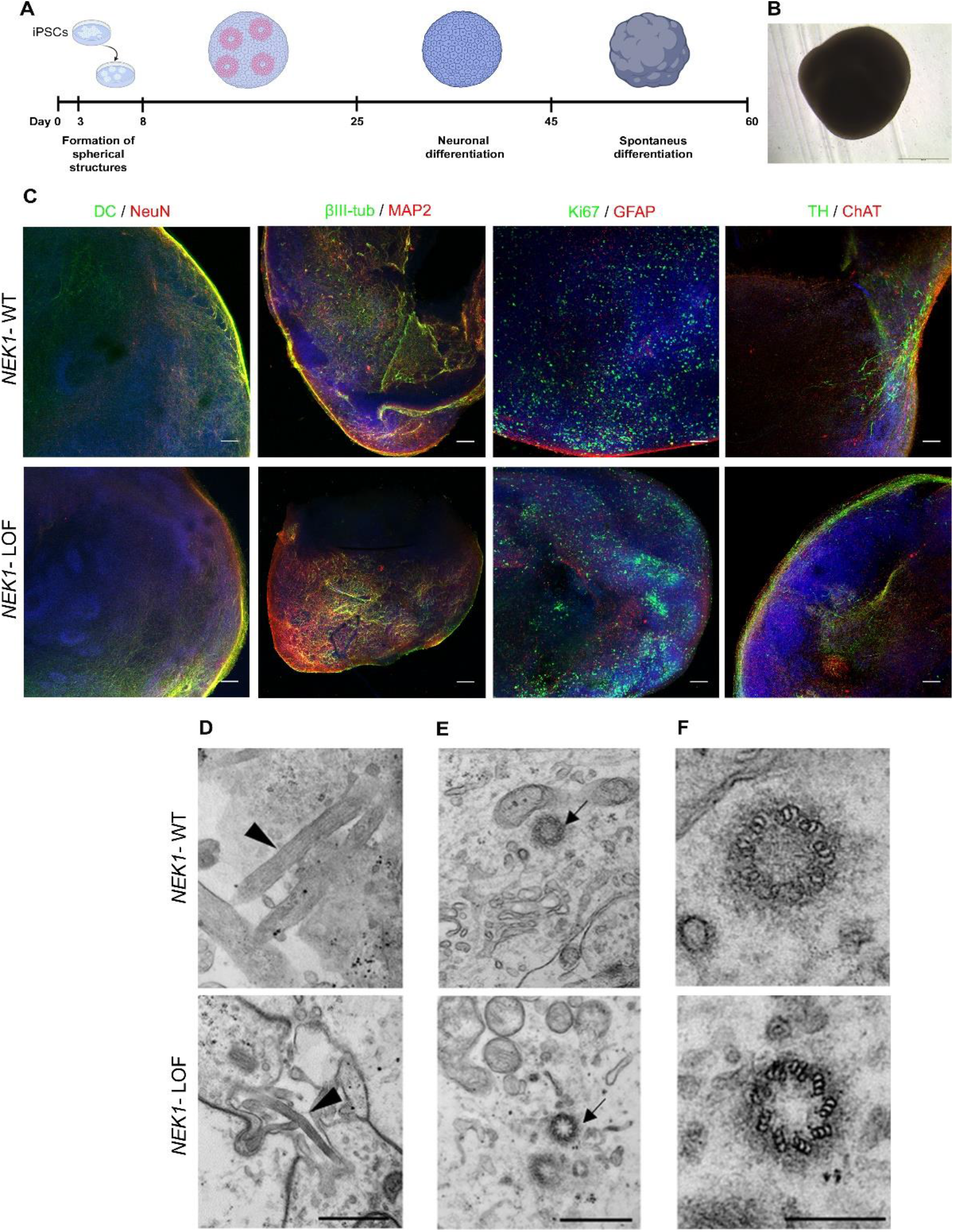
Analysis of primary cilia in human iPSC-derived brain organoids. (A) Scheme illustrating the differentiation steps for generating human brain organoids from iPSCs. (B) Morphology and size of a human brain organoid at day 60 of maturation. (C) Representative confocal images of neuronal markers βIII tubulin, Doublecortin (DC), Ki67 and Tyrosine Hydroxylase (TH) (all green) and MAP2, NeuN, GFAP and ChAT (all red) in *NEK1*-WT and *NEK1*-LOF iPSC-brain organoids. Scale bar 10 μm. Transmission electron microscopy of brain organoids. (D) Longitudinal sections (arrowhead) and (E) cross sections (arrow) of a primary cilium in *NEK1*-WT and *NEK1*-LOF iPSC-brain organoids and (F) enlarged cross section images shown in (E) with the detailed 9 + 0 architecture of the primary cilium at transition fiber. Scale bar 500 nm in (D) and (E); scale bar 250 nm in (F)

## DISCUSSION

NEK1 is a widely expressed 130 KDa serine/threonine kinase that plays crucial roles in various cellular processes, including cell cycle progression, DNA damage response, mitochondria integrity, nuclear import, microtubule organization and primary cilia formation (20).

In cilia, NEK1 regulates the organization of microtubules in the axoneme and/or basal body and may therefore control cilia length (21). *NEK1* LOF mutations in a recessive state cause STRD and SMDAX, two human ciliopathies characterized by skeletal dysplasia (7–9). Since heterozygous *NEK1* LOF mutations are instead associated with the motoneuron ALS disease, we investigated whether NEK1 haploinsufficiency impacts functionally on ciliogenesis by using human iPSC-derived motoneurons and brain organoids as 2D and 3D neural disease models.

Of interest, our data indicate that NEK1 protein deficiency and likely its reduced kinase activity, lead to defects in ciliogenesis with reduced primary cilium formation and thinner and shorter organelles in both ALS disease models of iPSC-MNs and iPSC-brain organoids. The use of iPSC-brain organoids certainly reinforces the reliability and robustness of our findings obtained in iPSC-MNs since iPSC-brain organoids allow to capture three-dimensional complexity and to more accurately represent human pathophysiology and non-cell autonomous mechanisms in ALS.

Our results define an association between *NEK1* LOF mutations and defects in neuronal primary cilia, but doesn’t establish causation. It’s unclear if primary cilia dysfunction might directly contribute to ALS pathogenesis or be a secondary effect, given the other important cellular processes which are also altered in condition of *NEK1* haploinsufficiency in human iPSC-MNs, including DNA damage repair (13) and nucleo-cytoplasm transport (14). Indeed, it was recently demonstrated that NEK1 binds and phosphorylates the tubulin subunit TUBA1B and contributes to regulate microtubules homeostasis and neurite growth in iPSC-MNs (14), strongly supporting the role of NEK1 also in the homeostasis of the microtubule-based primary cilium.

Noteworthy, primary cilia have already been reported to be shortened in both Alzheimer’s and Parkinson’s disorders, suggesting a possible mechanistic link between a defective cilium length and the neurodegeneration process (22). Therefore, understanding how cilia shortening can affect neural homeostasis in the context of *NEK1* haploinsufficiency in ALS disease will be crucial. Indeed, in neuronal cells primary cilia are important to regulate autophagy and proteostasis, brain energy homeostasis as well as mitochondrial functionality and senescence, well-recognized hallmarks of aging and neurodegenerative diseases (22). C21ORF2, a NEK1 protein interactor (23) and a genetic risk factor for ALS identified by a large genome-wide association study, (24) has also been recently associated to defective cilia formation and functioning (25). These functional data further reinforce the important role exerted by the NEK1-C21ORF2 axis in regulating ciliogenesis in ALS etiology, which certainly deserves further investigation.

Altogether our findings, by showing a link between *NEK1* LOF mutations and ciliogenesis in human 2D and 3D iPSC-derived neuronal disease models, suggest that primary cilia dysfunction may represent a still poorly considered pathomechanism in ALS, but a promising and novel target for addressing future therapeutic strategies for this uncurable disease.

## ABBREVIATIONS

ALS: Amyotrophic Lateral Sclerosis
NEK1: NIMA - related kinase1
WT: Wild type
2D: two dimensions
3D: three dimensions
LOF: Loss of function
SRTD: Short-rib thoracic dysplasia
SMDAX: Axial spondylometaphyseal dysplasia
iPSCs: Induced Pluripotent Stem Cell
EBs: Embryoid bodies
MNs: Motoneurons
TEM: Transmission Electron Microscopy

## DECLARATIONS

### Ethics approval and consent to partecipate

The study was approved by IRCCS Istituto Auxologico Italiano Research Ethics Board (title of the approved project: “iPSCs: un modello per lo studio delle interazioni tra cellule neurali nelle malattie neurodegenerative”; approval number: 2015_03_31_07; date of approval: 03/31/2015).

Skin biopsies and blood of healthy individuals and patients were obtained after written informed consent.

### Consent for pubblication

Not applicable.

### Author contributions

AR and MNS conceived and designed the study. Material preparation, data collection and analysis were performed by MNS, SI, VC, SS, CL, PP, AB, AS, SP, AQ and PB. The draft of the manuscript was written by MNS and AR. AR, PB and VS supervised the work. All authors read and approved the manuscript.

## Acknowledgements

We acknowledge Dr. Alessandra Sironi for the cytogenetic analyses and and Dr. Jan Hensen (University of Bonn, Germany) for his help with the CiliaQ plugin. SI, VC and SS are recipients of fellowships from the PhD program in “Experimental Medicine”, Università degli Studi di Milano. AR acknowledges “Aldo Ravelli Center for Neurotechnology and Experimental Brain Therapeutics”, Università degli Studi di Milano.

## Data availability statement

Original data are available upon reasonable request at Zenodo repository (doi: 10.5281/zenodo.10210708).

## SUPPLEMENTARY DATA

**Table 1.**
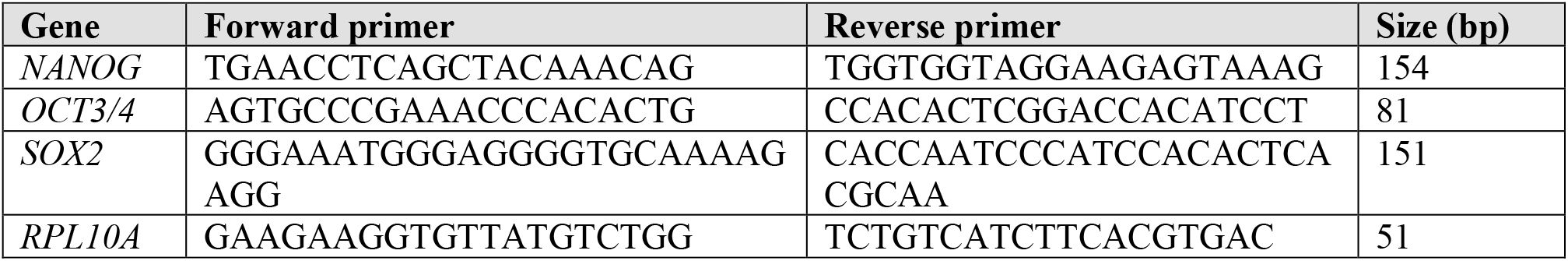
List of primer sequences for Real-time PCR.

**Table 2.**
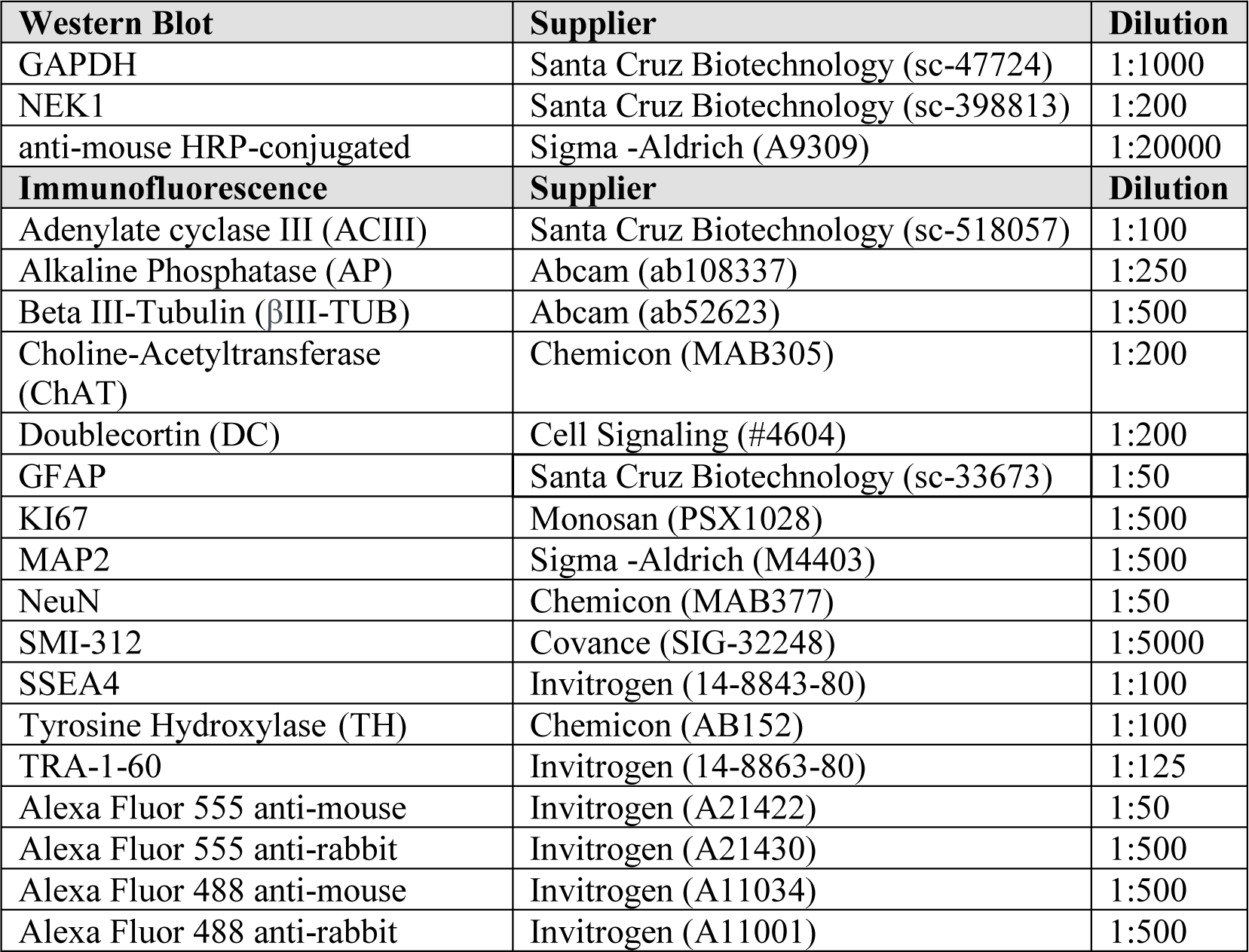
List of primary and secondary antibodies.

